# ProxECAT: Proxy External Controls Association Test. A new case-control gene region association test using allele frequencies from public controls

**DOI:** 10.1101/271007

**Authors:** AE Hendricks, S Billups, HNC Pike, IS Farooqi, E Zeggini, SA Santorico, I Barroso, J Dupuis

## Abstract

A primary goal of the recent investment in sequencing is to detect novel genetic associations in health and disease improving the development of treatments and playing a critical role in precision medicine. While this investment has resulted in an enormous number of sequenced genomes, individual studies of complex traits are often smaller and underpowered to detect rare variant genetic associations. Existing genetic resources such as the Exome Aggregation Consortium (>60,000 exomes) and the Genome Aggregation Database (^~^140,000 sequenced samples) could be used as controls in these studies. Fully utilizing these and other existing sequencing resources has the potential to increase power and could be especially useful in studies where resources to sequence additional samples are limited. However, to date, these large, publicly available genetic resources remain underutilized, or even misused, in large part due to the lack of statistical methods that can appropriately use this summary level data. We present a new method to incorporate external controls in case-control analysis called ProxECAT (Proxy External Controls Association Test). ProxECAT estimates enrichment of rare variants within a gene region using internally sequenced cases and external controls. We evaluated ProxECAT in simulations and empirical analyses of obesity cases using both low-depth of coverage (7x) whole-genome sequenced controls and ExAC as controls. We find that ProxECAT maintains the expected type I error rate with increased power as the number of external controls increases. With an accompanying R package, ProxECAT enables the use of publicly available allele frequencies as external controls in case-control analysis.

## Introduction

Recent investments have produced sequence data on millions of people with the number of sequenced individuals continuing to grow. Although large sequencing studies, such as the Trans-Omics for Precision Medicine (TopMed) through the National Heart, Lung, and Blood Institute, exist, most sequencing data is gathered and processed in much smaller units of hundreds to thousands of samples. This is especially true in the study of diseases that are not very common but still likely to have a complex or oligogenic genetic architecture. These silos of data mean that most rare-variant association studies of uncommon, complex diseases are underpowered. Zuk et al. suggest that sample sizes in the tens, and perhaps hundreds of thousands are required for adequate power^1^. In addition to increasing the sample size of future studies, fully leveraging existing sequencing resources could increase power considerably and could be vital in scenarios where resources to sequence more samples are limited.

Existing genetic resources such as the Exome Aggregation Consortium (ExAC; >60,000 exomes)^2^ and more recently, the Genome Aggregation Database (gnomAD; ^~^140,000 sequenced samples) have the potential to be used as controls in studies of complex diseases. However, to date,these large, publicly available genetic resources remain underutilized, or even misused^3^, in large part due to the lack of statistical methods that can appropriately use this summary level data in complex disease studies. In particular, there is a large potential for bias caused by differences in sequencing technology, processing, and read depth^3^.

Recently, Lee et al^4^ developed iECAT, a method to incorporate publicly available allele frequencies from controls into an existing, unbiased, but underpowered case-control analysis. They found that iECAT controls for bias while increasing power to detect association to a genetic region and can be applied to both single variant analysis and gene region analysis using a SKAT-O framework^5^. iECAT cannot be applied to very rare variants such as singletons or doubletons and requires a set of controls that were sequenced and variant-called in parallel to the cases (i.e. internal controls). Additionally, the type I error rate for iECAT increases as the size of the internal control sample set decreases relative to the internal cases. Thus, there is still the need for methods that can incorporate very rare variants and external controls without the explicit need for large internal control samples.

Here we present Proxy External Controls Association Test (ProxECAT), a method to estimate enrichment of rare variants within a gene region using internal cases and external controls. Our method addresses existing gaps such as using singleton and doubleton variants and requiring only external controls.

Rare-variant tests in a gene are often limited to variants predicted to have a functional effect on the protein, hence discarding non-functional variants. This can result in greater power^6;7^. The development of ProxECAT was motivated by the observation that these discarded variants can be used as a proxy for how well variants within a genetic region are sequenced and called within a sample. ProxECAT is both simple and fast, requiring only allele frequency information, and is thus well suited to use publicly available resources such as ExAC and gnomAD.

We evaluate ProxECAT in simulations, and empirical analysis of high depth of coverage (80x) whole-exome sequenced childhood obesity cases (N=927) using both low-depth of coverage (7x) whole-genome sequenced controls (N=3,621), and ExAC (N=33,370). Our method controls the type I error rate in simulations and yields the expected distribution of test statistics in real data settings. Given an accompanying R package, ProxECAT provides a robust and previously unavailable method to use publicly available allele frequencies as external controls in case-control analysis. This increases the utility of existing sequenced datasets to generate hypotheses and further research into the genetic basis of disease.

## Subjects and Methods

### Proxy External Controls Association Test

For a gene region-based test, we consider the following. Let Y denote the disease status, with Y = 1 and Y = 0 for internal case and external control status, respectively. We split the variants into those that are predicted to have a functional genetic impact and those that are not predicted to have a functional impact. We use the latter as the proxy variants. Let, 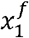 and 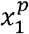 denote the counts of the functional and proxy rare variant alleles respectively for internal cases and 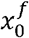 and 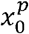 denote the counts of functional and proxy rare variant alleles respectively for external controls (Table 1).

**Table 1.** Data notation for internal case and external control samples for ProxECAT

|  |  | Predicted Functional Impact |  | Total |
| --- | --- | --- | --- | --- |
|  |  | Functional | Not Functional (Proxy) |  |
| Cases (Internal) | $Y = 1$ | $x_1^f$ | $x_1^p$ | $x_1$ |
| Controls (External) | $Y = 0$ | $x_0^f$ | $x_0^p$ | $x_0$ |
| Total | | $x^f$ | $x^p$ | |

We model the observed variant minor allele counts in Table 1 as a random sample from four independent Poisson distributions, i.e., 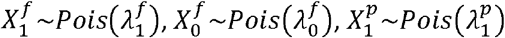, and 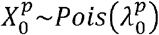. The derivation of the ProxECAT test statistic follows from the null hypothesis in Equation (1)

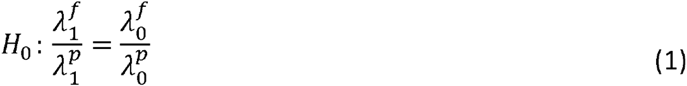

Using the method of Lagrange Multipliers and the constraint as defined by the null hypothesis, we find the maximum likelihood estimates (MLEs) of our parameters: 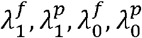. Details are in Supplemental Appendix A.

Our MLEs under the null hypothesis are:

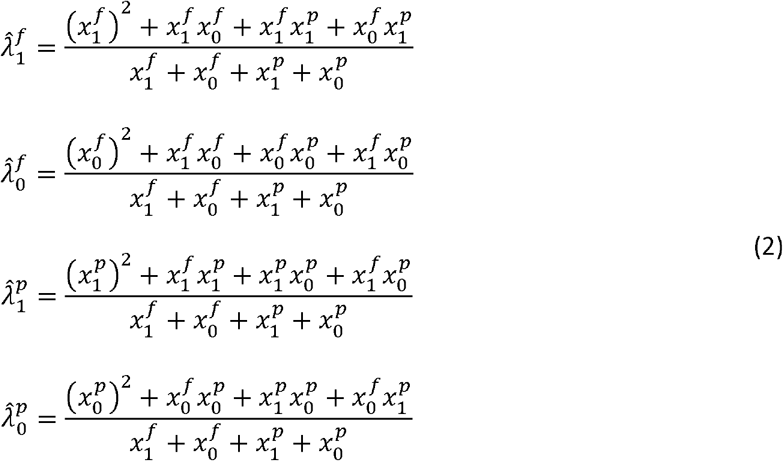

We use the parameter estimates in the likelihood for the constrained null hypothesis. The MLEs for the unconstrained alternative hypothesis parameters are the variant allele counts for each group 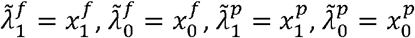. We then complete a likelihood ratio test (LRT) as the ratio of the constrained (null hypothesis) and unconstrained (alternative hypothesis)likelihoods, which, by Wilk’s theorem^8^ can be transformed to have a chi-squared distribution with 1-df.

### Extension to Incorporate Different Depths of Coverage

It has been shown that functional variants have a lower minor allele frequency (MAF) distribution compared to synonymous variants^9^. Further, high-depth of coverage sequencing will detect a higher number of variation at lower MAFs compared to low-depth of coverage sequencing^9; 10^. This results in high-depth of coverage sequencing detecting more functional variation relative to synonymous variation compared to low-depth of coverage sequencing. To allow for scenarios where sequencing coverage varies considerably between cases and controls, we weight the observed functional variant minor allele counts. Specifically, we divide the number of minor alleles for functional variants by the median ratio of the number of minor alleles for functional to synonymous variants within cases (*M*_1_) and within controls (*M*_0_) separately.

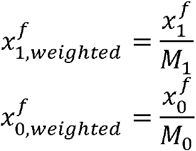

The weighted functional variant minor allele counts, 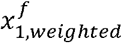 and 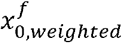, are used in place of the observed functional variant minor allele counts, 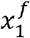 and 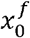, respectively to estimate the parameters in (2). This new test statistic is called ProxECAT-weighted.

### Extension to Negative Binomial

By assuming a Negative Binomial distribution for the number of minor alleles in a region instead of a Poisson distribution, we extend ProxECAT to incorporate possible over-dispersion. We model the Negative Binomial distribution with the mean, *λ*, and over-dispersion, *η*, parameters where the distribution approaches Poisson as *η* becomes large (Supplemental Figure 1).

### Software and Statistical Analysis

All tests were implemented using functions from our accompanying R package ProxECAT (https://github.com/hendriau/ProxECAT). Our primary test, which can model both ProxECAT and ProxECAT-weighted, was implemented with the *proxecat* function and our secondary test modeling over-dispersion was implemented using the *proxecat*.over function. We also implemented a **case-control LRT** to test for enrichment of rare, functional variant alleles in cases vs. controls and a **case-only LRT** similar to that performed by Zhi and Chen in 2012^11^. The case-only LRT tests for enrichment of rare alleles for functional variants in each gene of interest compared to the genome-wide average number of minor alleles per gene in cases only adjusting for the length of each gene. Unless otherwise specified, we assumed the data follow a Poisson distribution for all LRTs.

### Type I Error and Power Simulations

We simulated a variety of confounding scenarios. Case-control confounding represents systematic, genome-wide differences in the number of rare minor alleles between cases and controls. Gene confounding refers to a gene having a higher number of rare minor alleles than expected based on gene length. Gene confounding might occur due to several reasons including having a higher mutation rate, or poor annotation. The simulation scenarios and parameters are presented in Table 2 and Supplemental Table 1.

**Table 2.**
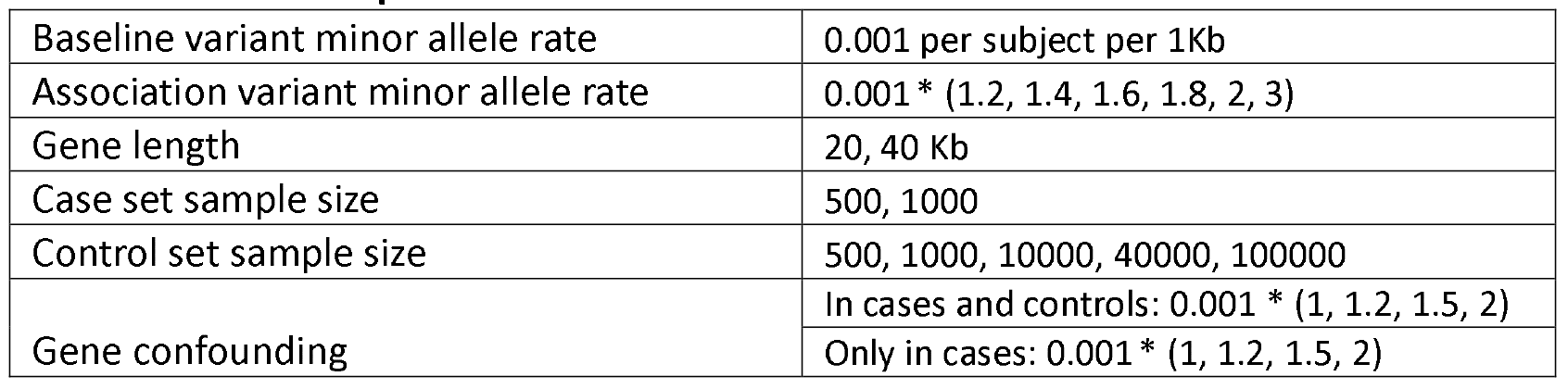
**Simulation parameters.**

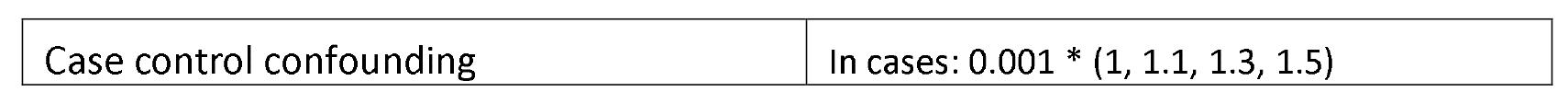

Within each case-control confounding simulation, we simulated 20,000 independent genes under four gene-disease association and gene confounding states. The four distinct gene states are: (1) association with case status and no gene confounding, (2) association with case status and gene confounding, (3) no association with case status and gene confounding, (4) no association with case status and no gene confounding. The number of rare minor alleles per gene was simulated under a Poisson distribution or an over-dispersed Poisson modeled using a Negative Binomial parameterization using the R functions *rpois* and *rnbinom*, respectively. The mu and size parameters in *rnbinom* represent the mean and over-dispersion, respectively.

### Assessing Fit of the Poisson Distribution

We assessed the validity of assuming a Poisson distribution for the number of minor alleles observed in a gene region. We simulated the number of each genotype group for each variant assuming Hardy-Weinberg Equilibrium and a Binomial distribution where p was the MAF. We varied the MAF (0.0001, 0.0005, 0.001, 0.005), the sample size (1000; 10,000), and the maximum number of variable variants within the gene region (5,10,20). We then assessed how closely the simulated distributions of the number of minor alleles observed per gene region matched a theoretical Poisson distribution where λ was the mean from each simulation scenario.

### UK10K SCOOP

Whole-exome sequenced (WES) cases are from the Severe Childhood Onset Obesity Project (SCOOP) cohort^6; 12^, which is a self-reported UK European subset of the Genetics of Obesity Study (GOOS). GOOS includes individuals with severe early-onset obesity body mass index (BMI) standard deviation score (SDS) > 3 and age at onset of obesity < 10 years. Leptin deficient individuals (identified by biochemical measurement) and those with mutations in the *MC4R* gene were excluded.

We used VerifyBamID (v1.0)^13^ and a threshold of ≥3% to identify contaminated samples. We computed principal components with the 1000Genomes Phase I integrated call set^9^ using EIGENSTRAT v4.2^14^ to identify non-Europeans, and pairwise identity by descent estimates from PLINK v1.07^15^ with a threshold of ≥0.125 to identify related individuals. Contaminated, non-European, and related samples were removed resulting in 927 SCOOP cases for analysis. Details about sequencing and variant calling for the SCOOP cases, as part of the UK10K exomes can be found elsewhere ^16^. All participants gave written informed consent and all methods were performed in accordance with the relevant laboratory/clinical guidelines and regulations.

### UK10K Cohort

The whole-genome sequenced (WGS) controls consist of the UK10K Cohort sample, comprised of two population cohorts: the Avon Longitudinal Study of Parents and Children (ALSPAC) and the TwinsUK study from the Department of Twin Research and Genetic Epidemiology at King’s College London (TwinsUK). We used allele frequency data for 3,621 individuals that passed sample QC as described elsewhere^17^.

### Exome Aggregate Consortium

We used allele frequency values for the N=33,370 non-Finnish European (NFE) group from the ExAC variant site dataset version 1.0 (http://exac.broadinstitute.org/downloads)^2^.

### Variant and Gene Filtering

To focus on rare or very rare variants, we limited to variants below a pre-specified MAF threshold in both cases and controls. We used MAF ≤ 1% in the SCOOP cases vs. UK10K cohort controls analysis and MAF ≤ 0.1% in the SCOOP vs. ExAC analysis. For the SCOOP cases vs. UK10K controls analysis, we also applied external filtering excluding variants with a MAF > 1% in at least one of the 1000Genomes five primary ancestry groups. Exclusion by 1000Genomes MAF was not possible when using ExAC as 1000Genomes sample are included in the ExAC genotype frequencies. We explored the distribution of test statistics over several thresholds for the minimum number of functional (*x*^*f*^) and proxy (*x*^*p*^) variants within each gene (5,10,20).

Analysis regions were limited to the intersection of respective target regions for SCOOP vs. UK10K Cohort and for SCOOP vs. ExAC. All variant annotation was applied using the GRCh37 human reference. The Ensembl Variant Effect Predictor (VEP, http://www.ensembl.org/info/docs/variation/vep/index.html^18^ v79 and v90.1) from Ensembl was used to add variant consequence annotations for SCOOP vs. UK10K Cohort and SCOOP vs. ExAC respectively. We defined functional variation using the following Sequence Ontology terms^19^ variant consequences: splice_donor_variant, splice_acceptor_variant, stop_gained, frameshift_variant, stop_lost, initiator_codon_variant, inframe_insertion, inframe_deletion, missense_variant, and protein_altering_variant. Variants were considered synonymous if they had the “synonymous_variant” flag.

### Assessing Results from Real Data Analysis

We used quantile-quantile plots (QQ-plots) to assess the resulting distribution of test statistics from the real data applications. Specifically, we looked at the middle of the distribution of test statistics as assessed by the lambda value (i.e. the median of the observed test statistic divided by the median of the expected test statistic) and the tail of the distribution of test statistics, which we assessed visually.

## Results

### Assessing Fit of the Poisson Distribution

In our comparison of the simulated number of rare minor alleles assuming a Binomial distribution to the theoretical Poisson distribution, no over dispersion was apparent. The sampling mean and variance of the simulated scenarios were similar across different sample sizes, MAFs, and number of minor alleles per gene (Supplemental Figures 2 and 3). When the expected number of minor alleles per gene was greater than 20, the Poisson approximation for the number of minor alleles started to look more continuous. In other words, as the expected number of variants per gene decreased, the Poisson approximation became more discrete and multimodal (Supplemental Figures 2 and 3). The theoretical distribution for the number of minor alleles per gene created from simulating genotypes for individual, independent variants from a Binomial distribution was more robust to discretization maintaining a mostly continuous distribution until the expected number of minor alleles per gene was equal to or less than four.

### Type I Error and Power Simulation Results

The case-control LRT (see Software and Statistical Analysis under Subjects and Methods) was robust to gene confounding scenarios maintaining the appropriate type I error rate, but had an increased type I error rate in the presence of case-control confounding. The case-only LRT maintained appropriate type I error rate in the presence of case-control confounding but was inflated in the presence of gene-confounding. The inflation in the type I error for the case-control LRT and the case-only LRT increased further when both gene and case-control confounding were present. This was especially true for the case-control LRT (Figure 1).

**Figure 1.**
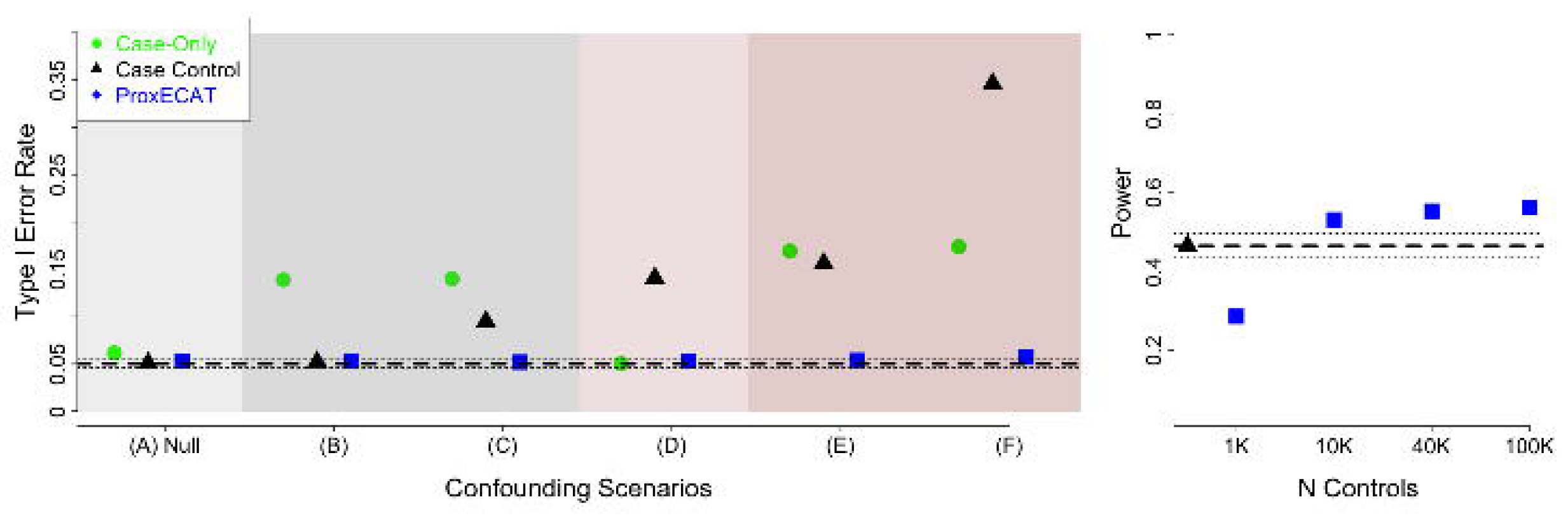
Type I error and power estimates for case-only LRT, case-control LRT, and ProxECAT over various confounding simulation scenarios. General simulation parameters: gene-length = 20Kb, baseline mutation rate = 0.001 per person per 1Kb. **Left Plot**: type I error rate for N_cases_=N_controls_=1000 and combinations of case-control confounding (mid level) and gene confounding (low level); dashed line represents expected type I error rate of 0.05 and dotted lines represent 95% confidence interval around the expected type I error rate. (A) Null simulation with no case-control or gene confounding bias; (B) gene-confounding; (C) gene-confounding only in cases; (D) case-control confounding; (E) case-control confounding and gene confounding; (F) case-control confounding and gene confounding only in cases. **Right Plot**: power for an effect size of 2 for case-control LRT (Ncases = 500; Ncontrols = 500) and ProxECAT (Ncases = 1000) and various external controls sample size. Dashed line is the case-control LRT power and dotted lines represent 95% confidence interval around the estimated power for case-control LRT.

Despite usually being within the 95% confidence interval for type I error, ProxECAT appeared to have a slight, but consistent inflation (Supplemental Table 2). This minor, but consistent inflation in the type I error rate can be addressed by using a more conservative significance threshold. We found that multiplying the significance level by 0.9 works well such that a 0.045 significance threshold maintains a 0.05 type I error rate, a 0.009 significance threshold maintains a 0.01 type I error rate, etc. Both the case-control LRT used here and ProxECAT assume a Poisson distribution and had inflated Type I Error rate in the presence of overdispersion (Supplemental Table 3). ProxECAT-over, which assumes a Negative Binomial distribution instead of a Poisson distribution, corrects for overdispersion in simulations when the overdispersion parameter is known and overdispersion is not too extreme (i.e. over-dispersion, η ≥ 5) (Supplemental Table 3).

Case-control LRT had higher power than ProxECAT under scenarios of no case-control confounding and given the same sample size (Supplemental Table 4). However, the power of ProxECAT increased as the sample size of the external control set increased eventually reaching higher power than the case-control LRT for the same number of internal sequences (Figure 1). This increase in power for ProxECAT is due, in part, to being able to sequence more cases with ProxECAT (N = 1000) than with a case-control LRT where sequencing resources need to be split between cases and controls (here Ncases = 500 and Ncontrols = 500). ProxECAT’s power increased while the type I error stayed the same under confounding scenarios where the number of functional variants in the cases increases (Supplemental Table 4).

### SCOOP Data Analysis

We evaluated ProxECAT using the SCOOP sample as cases and either the UK10K Cohort or ExAC NFE as controls. High-depth of coverage WES SCOOP cases vs. low-depth of coverage WGS UK10K Cohort controls had an inflated distribution of test statistics for the case-control LRT both at the center (lambda = 1.971) and in the tail of the distribution. While we did not observe inflation in the tail of the distribution for ProxECAT (Figure 2), there was a large inflation in the overall distribution of test statistics (lambda = 3.151). We observed a much higher ratio of the number of minor alleles in functional to synonymous variants per gene for the high-depth of coverage cases, median = 3.00, versus the low-depth of coverage controls, median = 1.89 (Table 3). ProxECAT-weighted, which adjusts for this systematic difference in sequencing coverage, resulted in a distribution of observed test statistics that more closely matches the expected distribution (lambda = 1.026, Figure 2).

**Figure 2.**
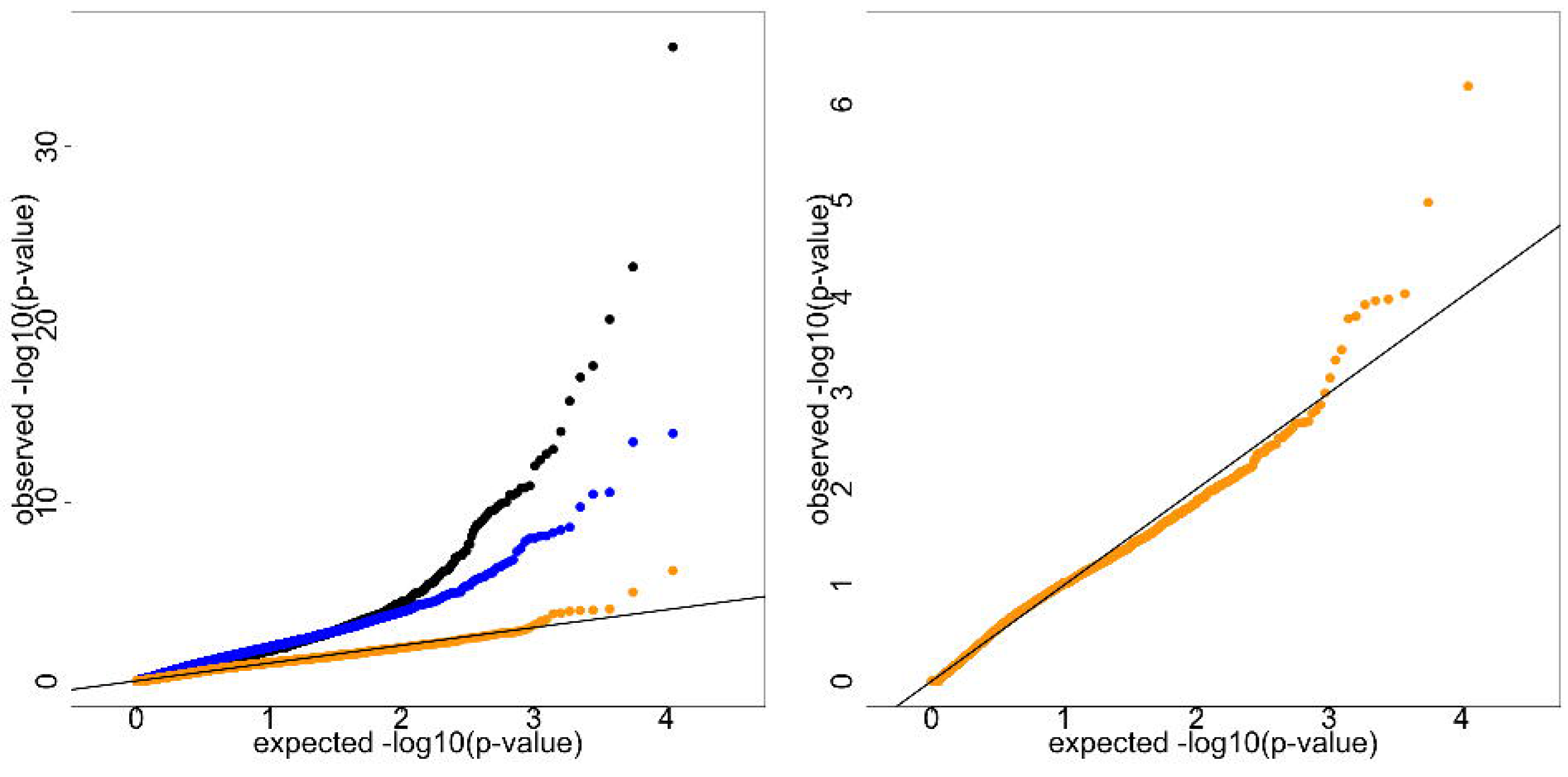
QQ plots for SCOOP cases vs. UK10K Cohort controls. Internal MAF < 0.01 in both cases and controls and number of variant minor alleles per gene ≥ 5. N genes = 11,051. ProxECAT (blue, lambda = 3.151, ProxECAT-weighted (orange, lambda = 1.026), case-control (black, lambda = 1.971). A) all tests, B) ProxECAT and ProxECAT-weighted only.

**Table 3.**
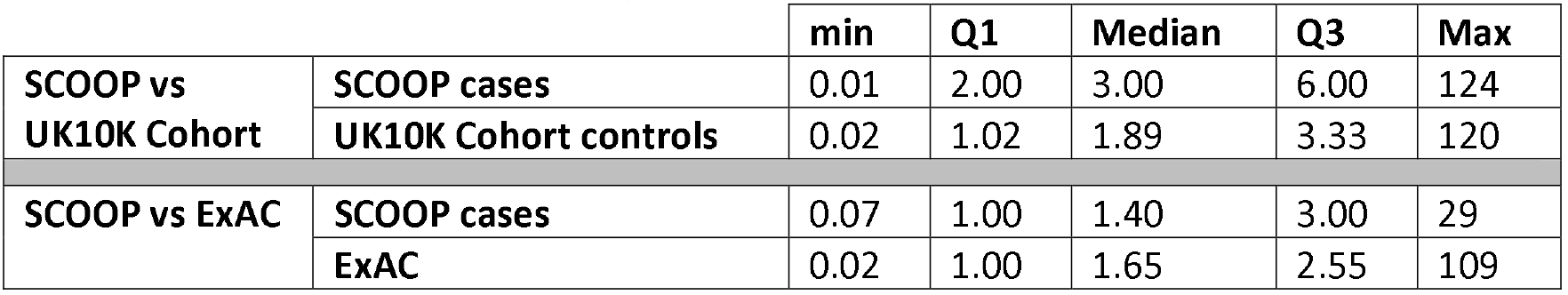
Genome-wide descriptive statistics for the ratio of the number of functional and synonymous variant minor alleles per gene in cases and controls

|  |  | min | Q1 | Median | Q3 | Max |
| --- | --- | --- | --- | --- | --- | --- |
| SCOOP vs UK10K Cohort | SCOOP cases | 0.01 | 2.00 | 3.00 | 6.00 | 124 |
|  | UK10K Cohort controls | 0.02 | 1.02 | 1.89 | 3.33 | 120 |
| SCOOP vs ExAC | SCOOP cases | 0.07 | 1.00 | 1.40 | 3.00 | 29 |
|  | ExAC | 0.02 | 1.00 | 1.65 | 2.55 | 109 |

A large strength of this method is the ability to use allele frequency data directly, rather than individual level allele calls. To assess the ability of this method to use publicly available allele frequency data, we used ExAC allele frequencies as controls for the SCOOP cases. The standard case-control LRT was inflated at both the median, lambda = 1.713, and tail (Figure 3) while our method maintained the expected distribution of test statistics. Because the depth of sequencing coverage is comparable and high for both SCOOP cases and ExAC controls, ProxECAT-weighted produced similar results to the standard, un-weighted test.

**Figure 3.**
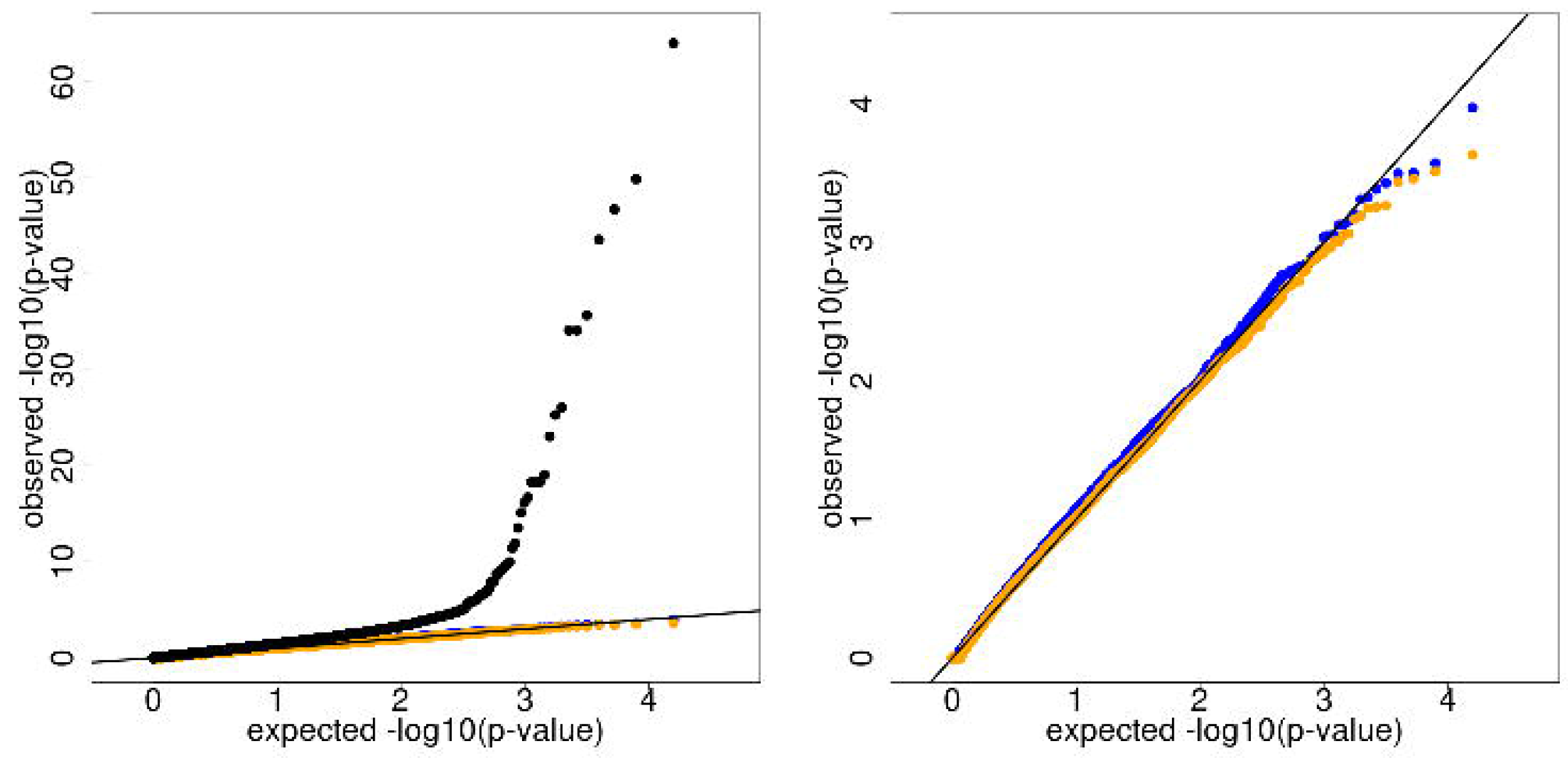
QQ plots for SCOOP cases vs. ExAC controls. Internal MAF < 0.001 in both cases and controls and number of variant minor alleles per gene ≥ 5. N genes = 15,863. ProxECAT (blue, lambda = 1.163, ProxECAT-weighted (orange, lambda = 1.069), case-control (black, lambda = 1.713) A) all tests, B) ProxECAT and ProxECAT-weighted only.

For both analyses, filtering to very rare variants was essential to avoid inflation in the distribution of observed test-statistics. This can be accomplished using moderate internal frequency filters and an external dataset such as 1000Genomes (MAF < 1%) as in the SCOOP vs UK Cohort analysis or using more stringent internal frequency filters (MAF < 0.1%) and no external dataset as in the SCOOP vs ExAC analysis.

Four genes, passing a 0.01 level of significance in both the SCOOP vs UK10K Cohort analysis and in the SCOOP vs ExAC analysis, are shown in Table 4. These results are putative novel obesity candidates meriting further replication. *MIB2* may be of particular interest as it is associated with decreased body weight in mice in the International Mouse Phenotyping Consortium (p-value = 7.49*10^−10^, http://www.mousephenotype.org/data/genes/MGI:2679684). Additional genes with the smallest p-values are found in Supplemental Tables 5-7.

**Table 4.**
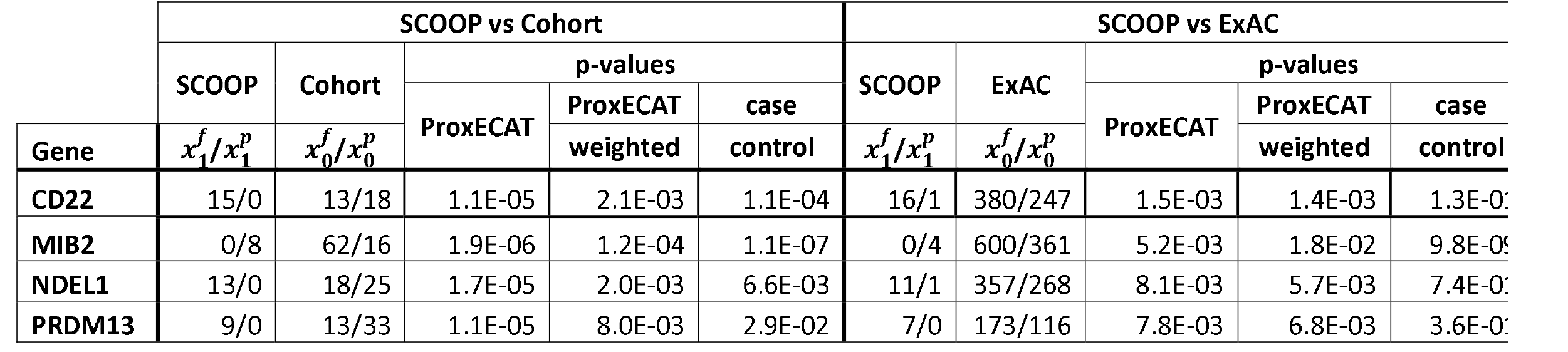
Gene-based results for genes with p-value < 0.01 in SCOOP vs. Cohort and SCOOP vs ExAC.

## Discussion

We propose a new method, ProxECAT, to test for enrichment of an accumulation of very rare variant alleles in a gene-region using publicly available external allele frequencies. ProxECAT only requires allele frequencies and uses exclusively external controls enabling the use of large, publicly available datasets such as ExAC and gnomAD. Analyses in simulations and using UK10K Cohort and ExAC as control sets for childhood obesity cases show that ProxECAT keeps the type I error rate and expected distribution of test statistics under control despite differences in sequencing technology and processing. Because ProxECAT uses external controls, additional resources can be devoted to sequencing cases. This results in greater power for ProxECAT compared to the case-control LRT test for the same number of internally sequenced individuals.

There are several limitations to the method proposed here. First, ProxECAT has a minor, but consistent inflation in the type I error rate. This limitation is easily addressed by using a more conservative significance threshold. Second, ProxECAT cannot currently include covariates such as sex, and ancestry. Thus, internal cases and external controls should be closely matched by ancestry and, as with any association study, findings will need independent replication preferably using a study where cases and controls are sequenced and processed in parallel. Third, the current approach does not enable internal controls to be analyzed along with external controls. While two analyses can be done in parallel and compared, it would be ideal to incorporate internal and external controls into the same statistical test. We are actively working on extensions to address these limitations.

It is important to highlight that research utilizing solely external controls is more susceptible to confounding due to known or unknown factors. Thus, any genes identified using ProxECAT or any method that uses only external controls should be carefully followed up in further validation, replication, and functional studies.

ProxECAT provides a robust approach to using allele frequencies from existing, publicly available sequencing data enabling case-control analysis when no or limited internal controls exist. ProxECAT uses the insight that readily available genomic information often discarded from analyses (here synonymous variation) can adjust for sizeable confounding due to differences in data generation. In the era of big data, we hope that both this insight and the ProxECAT method will enable additional genetic discoveries and will also motivate future methodological advancements in analyzing data across technologies and platforms.

## Acknowledgements

We are indebted to the patients and their families for their participation and to the physicians involved in the Genetics of Obesity Study (GOOS). This work was supported in part by the Burroughs Wellcome Fund (AEH) (Award #1015182) and National Institute of Health grant U01 DK078616 (JD). IB and EZ acknowledge funding from Wellcome (WT098051 and WT206194).

**R Package.** ProxECAT R package and functions are available on github: https://github.com/hendriau/ProxECAT

## Supplemental Figures

**Supplemental Figure 1.** Comparison of Poisson and Negative Binomial distributions for μ = 20.

**Supplemental Figure 2.** Comparison of the number of rare alleles in a gene region from a simulated variant level binomial distribution (black) and a theoretical Poisson distribution (red) for a sample size of 10,000. MAF = 0.0001, 0.0005, 0.001, 0.005; number of minor variant alleles within the gene region = 5, 10, 20.

**Supplemental Figure 3.** Comparison of the number of rare alleles in a gene region from a simulated variant level binomial distribution (black) and a theoretical Poisson distribution (red) for a sample size of 1,000. MAF = 0.0001, 0.0005, 0.001, 0.005; number of variants within the gene region = 5, 10, 20.

### Supplemental Tables

**Supplemental Table 1.** Gene confounding and confounding simulation design. Darker shading indicates a higher level of gene confounding. Solid shading indicates gene confounding in both cases and controls. Stripped shading indicates gene confounding in only cases.

**Supplemental Table 2.** Type I Error over all simulation scenarios.

**Supplemental Table 3.** Type I Error for over-dispersed simulations. Gene length = 20Kb, Ncases = 1000, Ncontrols = 1000, no confounding.

**Supplemental Table 4.** Power over all simulation scenarios.

**Supplemental Table 5.** Top 100 results for SCOOP vs. Cohort ordered by ProxECAT-weighted p-value.

**Supplemental Table 6.** Top 100 results for SCOOP vs. ExAC ordered by ProxECAT p-value.

**Supplemental Table 7.** Results with p-value < 0.05 for both SCOOP vs. Cohort and SCOOP vs. ExAC.

### Supplemental Information

**Supplemental Appendix A.** Derivation of ProxECAT.

